# Coral cover surveys corroborate predictions on reef adaptive potential to thermal stress

**DOI:** 10.1101/2020.06.05.136523

**Authors:** Oliver Selmoni, Gaël Lecellier, Laurent Vigliola, Véronique Berteaux-Lecellier, Stéphane Joost

## Abstract

As anomalous heat waves are causing the widespread decline of coral reefs worldwide, there is an urgent need to identify coral populations tolerant to thermal stress. Heat stress adaptive potential is the degree of tolerance expected from evolutionary processes and, for a given reef, depends on the arrival of propagules from reefs exposed to recurrent thermal stress. For this reason, assessing spatial patterns of thermal adaptation and reef connectivity is of paramount importance to inform conservation strategies.

In this work, we applied a seascape genomics framework to characterize the spatial patterns of thermal adaptation and connectivity for coral reefs of New Caledonia (Southern Pacific). In this approach, remote sensing of seascape conditions was combined with genomic data from three coral species. For every reef of the region, we computed a probability of heat stress adaptation, and two indices forecasting inbound and outbound connectivity. We then compared our indicators to field survey data, and observed that decrease of coral cover after heat stress was lower at reefs predicted with high probability of adaptation and inbound connectivity. Last, we discussed how these indicators can be used to inform local conservation strategies and preserve the adaptive potential of New Caledonian reefs.

## Introduction

Coral bleaching is one of the main causes of severe declines of coral reefs around the world^1-3^. This phenomenon is mainly caused by anomalous heat waves leading to the death of hard-skeleton corals, which are the cornerstone of reefs^2^. Over the last 30 years mass coral bleaching events repeatedly struck worldwide, causing losses of local coral cover up to 50%^1,3^. In the coming years, bleaching conditions are expected to occur more frequently and to become persistent by 2050^4^. As up to one third of marine wildlife depends on coral reef for survival and at least 500 million people livelihoods worldwide^5^, there is an urgent need to define new strategies to improve the preservation of these ecosystems^6^.

Recent research reported reefs that rebounded from repeated heat stress and showed an increased thermal resistance^7–11^. Adaptation of corals against heat stress might explain such observations^12,13^. Under this view, identifying adapted coral populations is of paramount importance, as conservation strategies might be established to protect reefs hosting these corals from local stressors (e.g. via marine protected areas, MPAs)^14^. Furthermore, adapted corals could be of use in reef restoration plans and repopulate damaged reefs^15^. The adaptive potential of corals at a given reef depends on the arrival of propagules from reefs exposed to recurrent thermal stress^16,17^. This is why characterizing spatial patterns of thermal adaptation and reef connectivity is crucial to empower the conservation of the adaptive potential of corals^16,17^.

Seascape genomics is a powerful method to evaluate spatial patterns of environmental variation and connectivity^17,18^. This method relies on a thorough analysis of environmental conditions around reefs using satellite data. Daily records of surface temperature are remotely sensed using satellites, and processed to compute indicators of thermal patterns associated with bleaching events^17,19,20^. Corals exposed to different thermal patterns are then sampled and genotyped to identify genetic variants correlated with these indicators^17,18^. The association between genetic variants and a given indicator defines a model of adaptation that can be used to predict the probability of adaptation, based on the value of the indicator itself^17,21^. In addition, by remote sensing sea current movements, it is possible to draw a connectivity map between every reef within an area of interest. This can be done using spatial graphs that resume multi-generational dispersal matching spatial patterns of genetic diversity in a given species^22^. This approach results in indices of connectivity defining, for a reef of interest, the predisposition in sending (outbound connectivity) and receiving (inbound connectivity) propagules to/from neighboring reefs^17^.

In this study, we predicted spatial patterns of heat stress adaptation and connectivity for over 1000 km of coral reefs of New Caledonia, in the Southern Pacific (Fig. 1). The study area encompassed the barrier reef surrounding Grande Terre, the main islands of the Archipelago, as well as the intermediary and fringing enclosed in the lagoon. We also considered reefs surrounding the Loyalty Islands (Ouvéa, Lifou and Maré) and the Astrolabe (east of Grande Terre) and those in the Entrecasteaux and Petri atolls (north of Grande Terre). We first used remote sensing data to (1) evaluate the thermal variability of the study area and (2) estimate patterns of sea current connectivity between reefs. Next, we employed genomic data from a seascape genomics study on three coral species of the region^23^ in order to (1) compute the probability of adaptation to heat stress across the whole region, and (2) verify whether predicted reef connectivity matched genetic correlation between corals. Last, we compared our predictions with field surveys of living coral cover recorded by the New Caledonian observational network of coral reef (RORC)^24^. Our results suggest that negative effects of recent heat stress on coral cover are mitigated at reefs predicted with high probability of heat stress adaptation and inbound connectivity. We then discuss the conservation status of reefs around New Caledonia, and assess how conservation indices of probability of adaptation and connectivity can be used to protect the adaptive potential of corals of the region.

**Figure 1.**
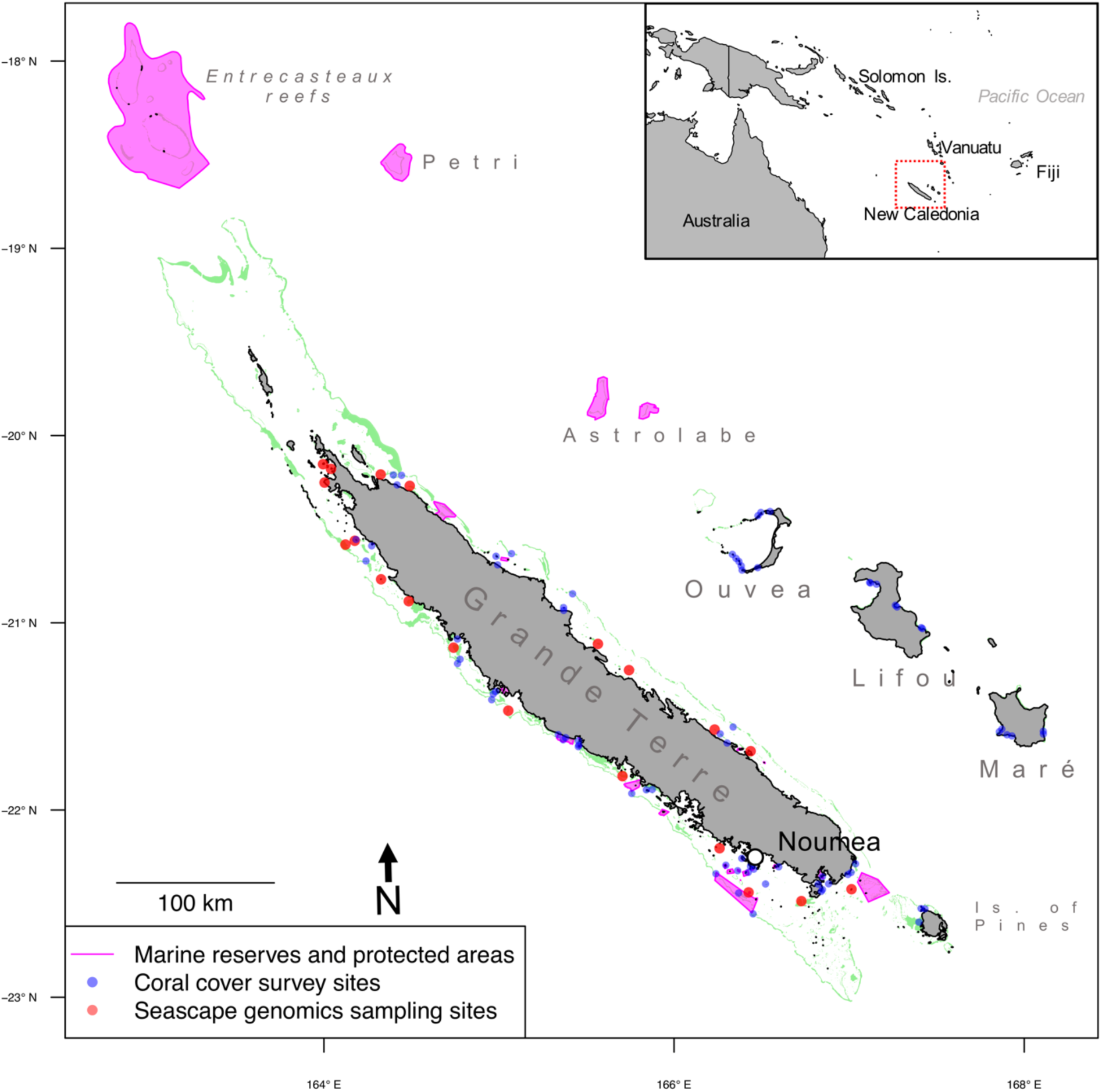
Reef system of New Caledonia. Coral reefs are highlighted in green. The blue dots correspond to sites of coral cover survey of the New Caledonian observational network of coral reef^24^. The red dots correspond to the sampling locations of coral specimen for the seascape genomics study that provide genetic data in the present study^23^. Sea regions highlighted in purple correspond to the marine reserves and protected areas as catalogued by the French agency for MPAs (http://www.aires-marines.fr/).

## Results

### Heat stress and probability of adaptation

The remote sensing data of sea surface temperature were processed to calculate the frequency of bleaching alert conditions (BAF_overall_) across the reef system of New Caledonia (Fig. 2a). BAF_overall_ was higher in reefs on the western coast of Grande Terre (average BAF: 0.16±0.04) than in those on the eastern coast (0.08±0.03). Reefs in Lifou, Maré and Isle of Pines displayed BAF_overall_ values comparable to those on the eastern coast of Grande Terre (0.09±0.03, 0.10±0.02 and 0.11±0.01, respectively), while in Ouvéa and Entrecasteaux reefs the BAF_overall_ values (0.15±0.01 and 0.12±0.01, respectively) were closer to the values observed on the western coast.

**Figure 2.**
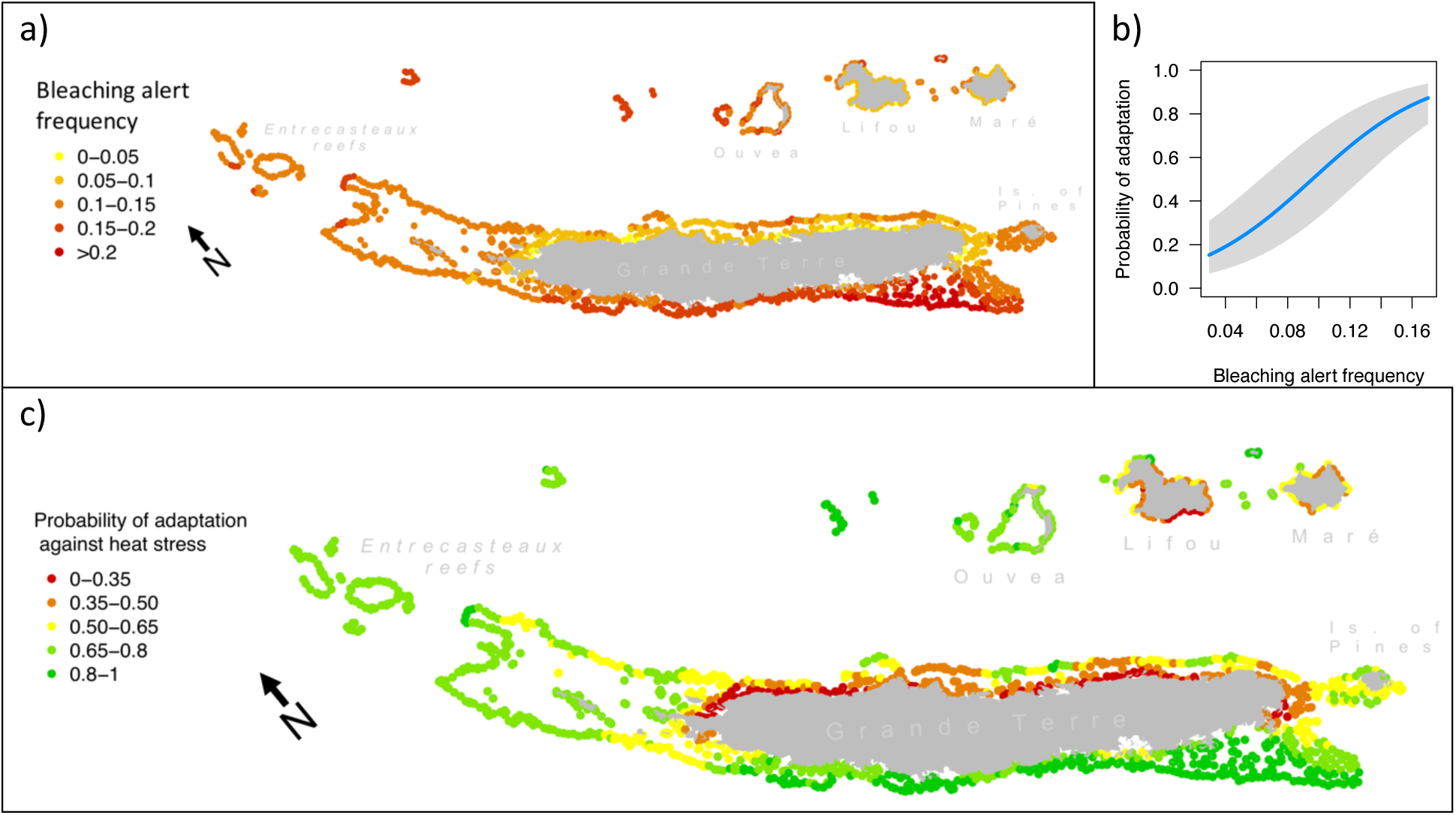
Bleaching alert frequency and probability of heat stress adaptation. In (a), bleaching alert frequency (BAF_overall_) is displayed for each reef of New Caledonia. This value is derived from remote sensing data of sea surface temperature, and describes the frequency of cumulated heat stress conditions that can lead to bleaching. In (b), a logistic model of heat stress adaptation is shown. This model is based on the frequencies of potentially adaptive genotypes of three coral species of New Caledonia^23^. The plot displays the probability of adaptation to heat stress as a logistic function of BAF_overall_ (blue line, with the grey band showing the 95% interval of confidence). The model shown in (b) was used to translate BAF_overall_ displayed in (a) in the probability of adaptation (PA_HEAT_) against heat stress. The map in (c) displays PA_HEAT_ for every reef of New Caledonia.

Previous seascape genomics analyses on three corals of the region (*Acropora millepora, Pocillopora damicornis and Pocillopora acuta*) revealed the presence of multiple genetic variants (32 in total) potentially implicated in heat stress resistance^23^. We employed this data to construct a logistic model of heat stress adaptation defining the probability of presence of potentially adaptive variants (PA_HEAT_) as a function of BAF_overall_ (Fig. 2b). This model was then used to produce a map of predicted PA_HEAT_ values for the whole region (Fig. 2c). It revealed accentuated differences compared with BAF_overall_ patterns, with PA_HEAT_ generally above 0.65 in reefs on the western coast of Grande Terre, Isle of Pines, Entrecasteaux and Ouvéa. In contrast, values below 0.35 were observed at reefs located along the east coast of Grande Terre, in Lifou and Maré.

### Reef connectivity and genetic correlation between corals

Remote sensing of sea current was used to compute a spatial graph of seascape connectivity predicting cost distances between reefs of New Caledonia. By using generalized linear mixed models (GLMMs) regression, we investigated whether such predictions on reef connectivity were representative proxies of the population structures of corals of the region. In three studied species (*A. millepora, P. damicornis and P. acuta*), we found that the genetic correlations between corals were significantly associated with the seascape cost distances separating the reefs where corals were sampled (*A. millepora*: p=1.46e-06; *P. damicornis*: p=1.63e-10 and *P*. acuta: p=6.7e-05; Fig. 3). This relationship was more stressed in the two *Pocillopora* species (regression coefficient for *P. damicornis*: *β*=-8.7E-05+ 1.4E-05; for *P. acuta*: *β*=-3.1E-05 ± 7.8E-06) than in *A. millepora* (*β*=-7.9E-06 ± 1.6E-06). The GLMMs accounted for the ancestral distance between pairs of individuals (*i*.*e*. the difference in admixture from ancestral populations) which was found significantly associated to genetic correlations between corals in all the three species (p∼0; Fig. S1).

**Figure 3.**
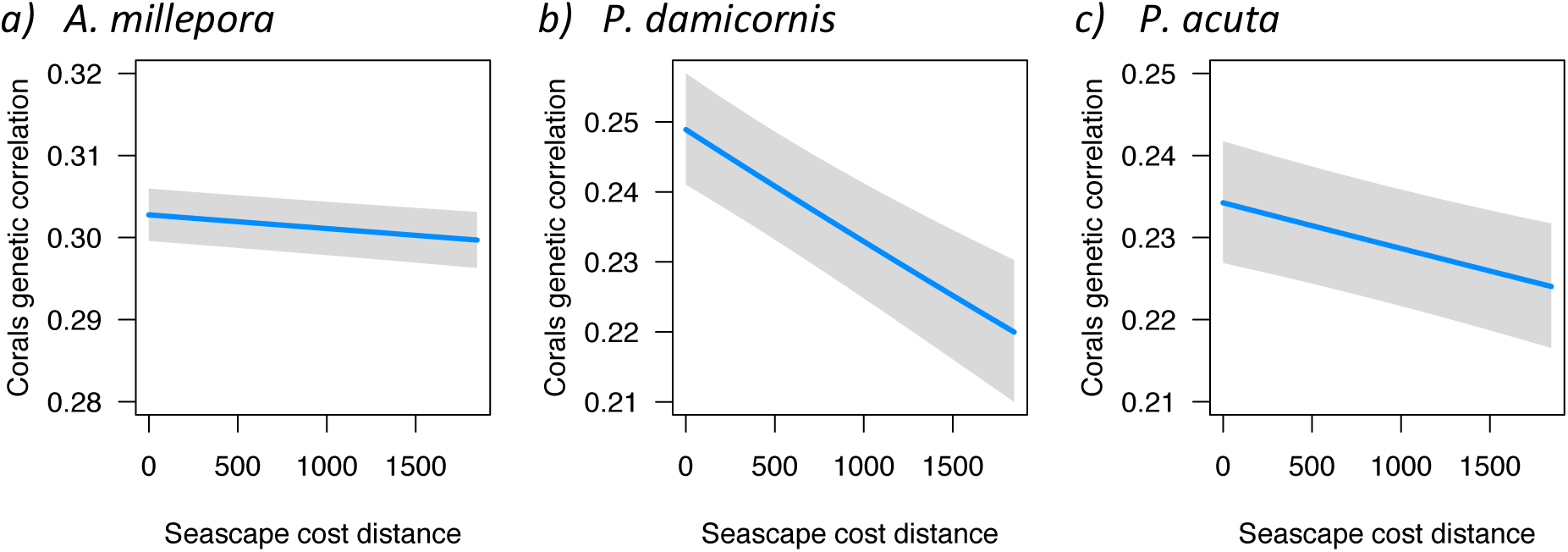
Seascape cost distance and genetic correlation between corals. The three plots display genetic correlations between pairs of corals sampled in New Caledonia as a function of the cost distance separating reefs where corals were sampled (blue line, with the grey band showing the 95% interval of confidence). Genetic correlations were computed as the correlation of single-nucleotide-polymorphisms, while seascape cost distance was predicted through seascape connectivity graphs. Each plot displays this association for a different species (a: *Acropora millepora*, b: *Pocillopora damicornis*, c: *Pocillopora acuta*).

### Reef connectivity indices

The seascape connectivity graph was used in the calculation of two indices describing the dispersal characteristics of every reef of New Caledonia (Outbound Connectivity Index, OCI, Fig. 4a; Inbound Connectivity Index, ICI, Fig. 4b). Both indices are expressed in km^2^, as they represent the area of the reefs neighboring a reef of interest. In OCI, neighboring reefs are those potentially receiving propagules from the reef of interest, while in ICI neighboring reefs are those potentially sending propagules towards the reef of interest.

**Figure 4.**
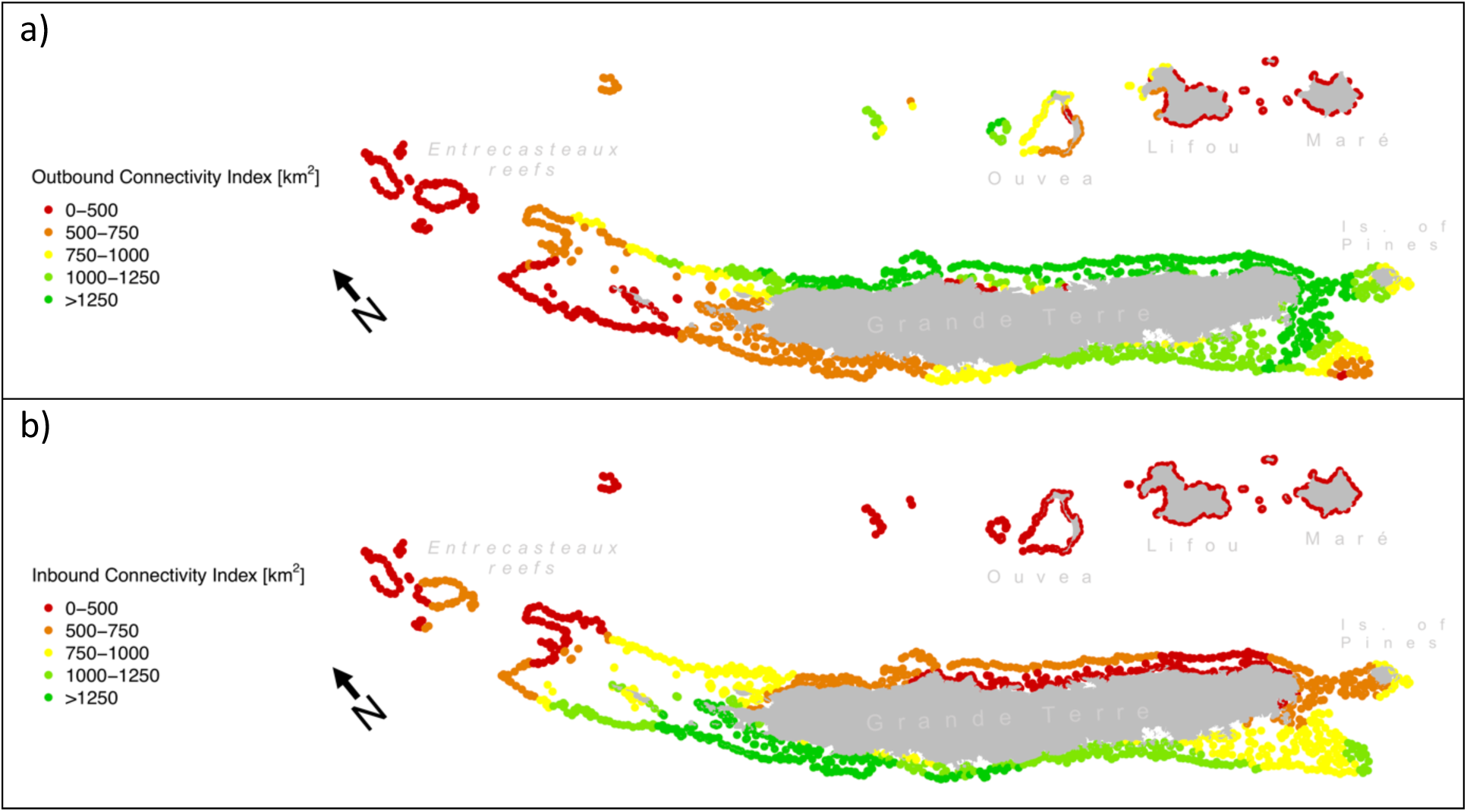
Connectivitiy indices. Two connectvitiy indices based on sea current data are shown for every reef of New Caledonia. In a), the Outbound Connectivity Index (OCI) describes the predisposition in sending dispersal to neighboring reefs. In b), the Inbound Connectivity Index (ICI) summarizes the predisposition in receiving propagules from neighboring reefs. Both indices are given in km^2^, as this represents the total surface of neighboring reefs.

Reefs that are more distant to Grande Terre (Entrecasteaux, Lifou, Maré and Ouvéa) had lower OCI (average OCI: 202±35 km^2^, 410±270 km^2^, 210±66 km^2^, 864±254 km^2^, respectively; Fig. 4a) than reefs surrounding Grande Terre. Reefs surrounding Grande Terre showed highest values on the southern reefs of the eastern coast (1929±300 km^2^), while lower values were predicted for the rest of the eastern coast (1377±435 km^2^) and the southern part of the western coast (1119±82 km^2^). OCI was lower at reefs located at the northern extremity of Grande Terre (632±244 km^2^).

Like with OCI, ICI was lower at reefs furthest from Grande Terre (Entrecasteaux, Ouvéa, Lifou, Maré; average ICI of 460±93 km^2^, 177±7 km^2^, 97±30 km^2^, 111±6 km^2^, respectively; Fig. 4b). ICI at reefs surrounding Grande Terre displayed a net contrast between the east and west coasts, where ICI was lower on the east (498±113 km^2^) than the west (1287±407 km^2^).

### Coral cover analysis

Underwater surveys of New Caledonian reefs were analyzed to characterize the association of living coral cover with recent thermal stress (BAF_previous year_), probability of heat stress adaptation (PA_HEAT_) and connectivity indices (ICI and OCI; Fig. 5). We first investigated the association between coral cover and individual explanatory variables using single fixed effect GLMMs (Fig. 5a-d). We found that coral cover was significantly associated with BAF_previous year_ (p=0.02), and that this association was of negative sign (*β*=-0.06±0.03; Fig. 5a). In contrast, none of the other univariate models resulted in a significant association with coral cover (PA_HEAT_: p=0.93, Fig. 5b; OCI: p=0.46, Fig. 5c; ICI: p=0.41, Fig. 5d). The Akaike Information Criterion (AIC) suggested a higher quality-of-fit for the model employing BAF_previous year_ as explanatory variable (AIC=-883), compared with the other univariate models (PA_HEAT_: AIC=-878; OCI: AIC=-879, ICI: AIC=-879).

**Figure 5.**
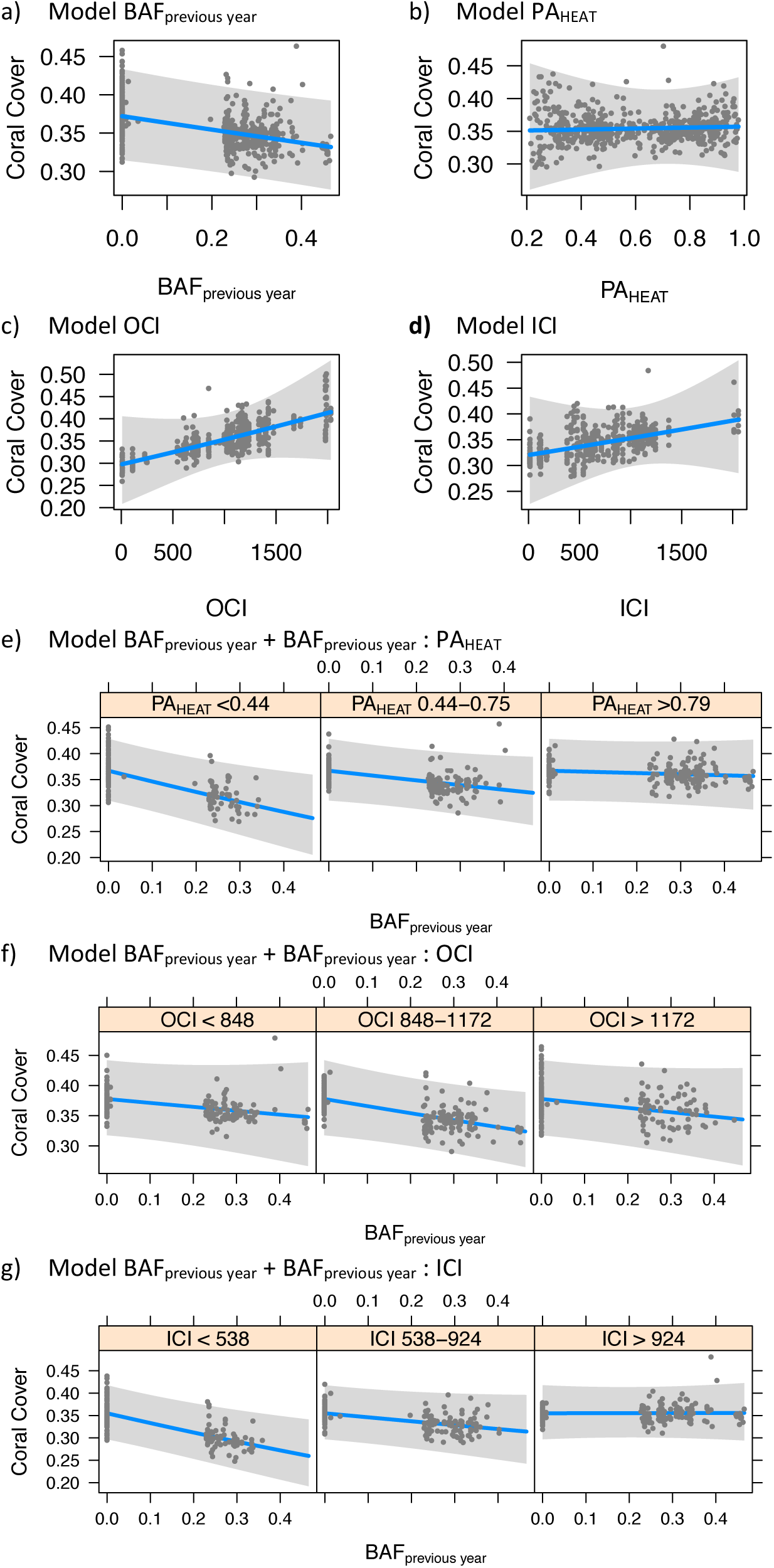
Coral cover association analysis. The plots display the association of coral cover rates (blue line, with the grey band showing the 95% interval of confidence) with recent thermal stress (BAF_previous year_), probability of heat stress adaptation (PA_HEAT_) and connectivity indices (inbound connectivity index, ICI, and outbound connectivity index, OCI). In plots (a) to (d), the association with coral cover rates is shown for each explanatory variable alone (a: BAF_previous year_, b: PA_HEAT_, c: OCI, d: ICI). In the remaining plots, the association between coral cover and BAF_previous year_ and is showed across different ranges PA_HEAT_ (e), OCI (f) and ICI (g).

We then investigated whether the negative association between coral cover and BAF_previous year_ varied under different values of PA_HEAT_, OCI or ICI. This analysis employed three bivariate GLMM setting as fixed effects BAF_previous year_ and the interaction between BAF_previous year_ and each of the three other explanatory variables (PA_HEAT_, OCI, ICI; Fig. 5e-g). In comparison to all the univariate models, those accounting for the interaction of BAF_previous year_ with PA_HEAT_ and ICI resulted in a higher quality-of-fit (AIC=-886 and AIC=-888, respectively). In both cases, the effect of BAF_previous year_ was significant (p<0.01) and of negative sign, whereas the effect of the interaction was also significant but of positive sign (for the interaction with PA_HEAT_: *β*=+0.05±0.02, p=0.03; with ICI: *β*=+0.07±0.03, p=0.01; Fig. 5e-f). In contrast, the bivariate model incorporating OCI had a quality-of-fit comparable to univariate models (AIC=-883), and showed no significant association in interaction with BAF_previous year_ (Fig. 5g).

## Discussion

### Local divergences in conservation indices

The metrics computed in this study stressed the strong asymmetry, in terms of both probability of heat stress adaptation (PA_HEAT_) and connectivity (inbound connectivity index, ICI; outbound connectivity index; OCI), between reefs on the two coasts of Grande Terre (Fig. 2a, Fig.4). The climatic differences between the two coasts are modulated by the mountain range covering Grande Terre, and water conditions inside the lagoon reflect the combination of these differences coupled with oceanic influences^25^. For example, the southern part of the west coast of Grande Terre is subjected to coastal upwelling, a seasonal phenomenon bringing cold water to the surface^26^. While logic would suggest that cold water alleviates heat stress, research on the Great Barrier Reef in Australia showed that intense upwelling is followed by severe heat stress, and consequent coral bleaching^27^. While it is unknown whether this same effect occurs on the south-western coast of Grande Terre, this region does enclose the reefs that are predicted to experience the highest frequency of bleaching conditions across New Caledonia, and consequently to host corals with the highest PA_HEAT_ (Fig. 2).

Asymmetrical spatial patterns between the coasts of Grande Terre were also predicted for connectivity (Fig. 4), and this matched the genetic population structure of corals of the region (Fig. 3). In this work, we estimated connectivity using a straightforward approach, conceived to be reproduceable on any reef system around the world but that might lead to local inaccuracies^17^. However, our predictions were generally consistent with previous work that characterized the regional water circulation around New Caledonia using more sophisticated methods (i.e. combining oceanographic models, in situ measurements and shipboard detectors of sea currents)^28^. For instance, we observed a higher inbound connectivity index (ICI) on the west coast of Grande Terre (Fig. 4b), and a higher outbound connectivity index (OCI) on the east coast (Fig. 4a). This west-oriented connectivity was expected because of the South Equatorial Current crossing the archipelago in this direction^28^. This current bifurcates at the encounter of the New Caledonian shelf into 1) a weak and transient south-east oriented current between the Loyalty Islands and Grande Terre, and 2) a strong north-west oriented current flowing north of the Loyalty Islands^26,28,29^. This bifurcation explains the lower OCI observed in Lifou and Maré, compared with Ouvéa and the Astrolabe atolls. Last, the water circulation inside the lagoon follows the north-west orientation of trade winds^26^, resulting in higher OCI in the south and higher ICI in the north.

Predictions of reef connectivity and PA_HEAT_ varied considerably across the different regions of the study area (Fig. 2, 4), and conservation planning should account for these regional peculiarities^14,30^. In table 1, we interpret the local divergences in values of PA_HEAT_, ICI and OCI under a conservation perspective.

**Table 1.**
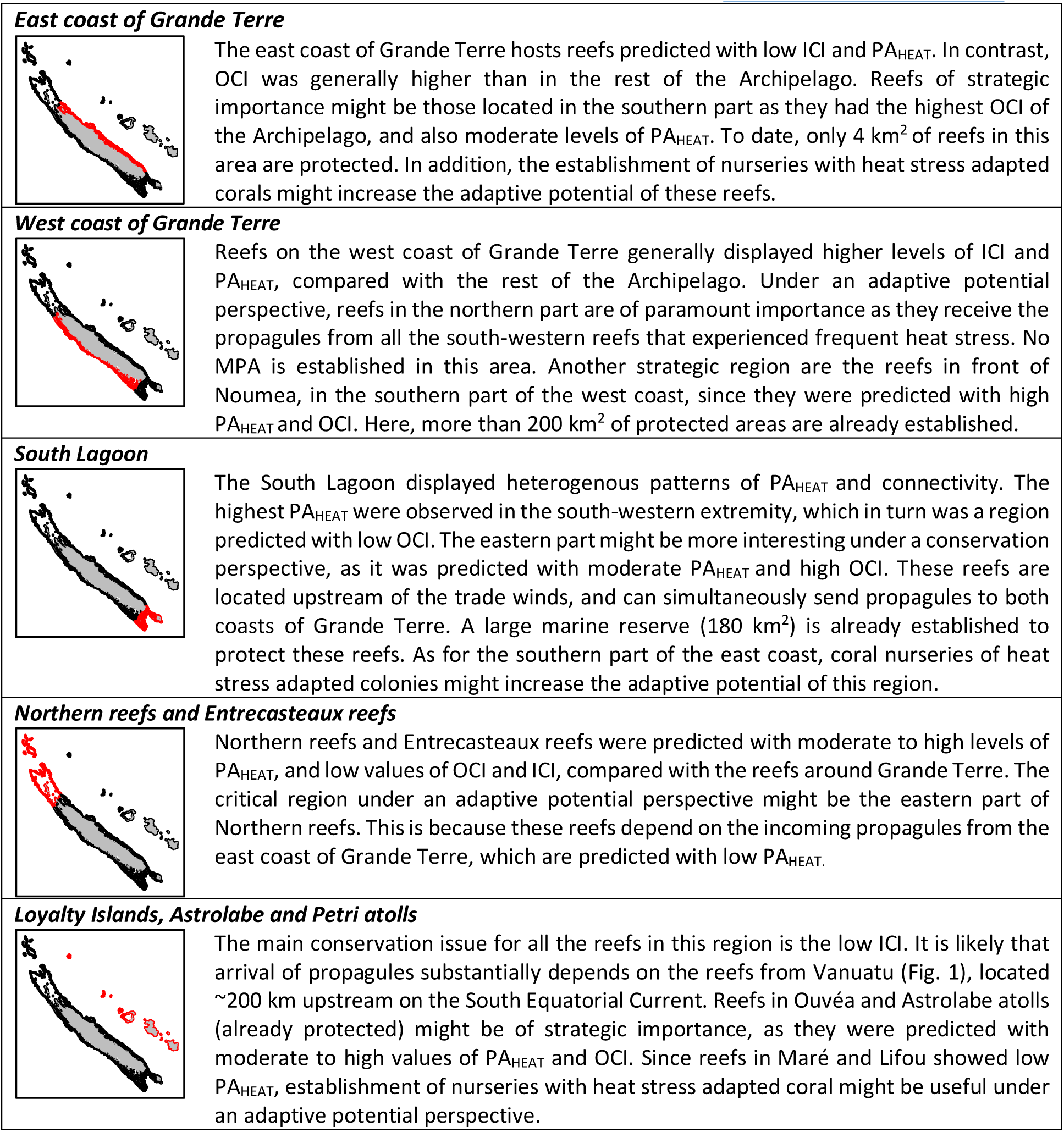
Implications for reef conservation in New Caledonia. The table describes the implications for reef conservation of the probability of heat stress adaptation (PA_HEAT_), the outbound and inbound connectivity indices (OCI, ICI) predicted for different regions of the New Caledonia reef system. Information on the existing marine protected areas were retrieved from the French agency for MPAs (http://www.aires-marines.fr/).

|  |  |
| --- | --- |
| **East coast of Grande Terre** | The east coast of Grande Terre hosts reefs predicted with low ICI and $PA_{HEAT}$. In contrast, OCI was generally higher than in the rest of the Archipelago. Reefs of strategic importance might be those located in the southern part as they had the highest OCI of the Archipelago, and also moderate levels of $PA_{HEAT}$. To date, only 4 km<sup>2</sup> of reefs in this area are protected. In addition, the establishment of nurseries with heat stress adapted corals might increase the adaptive potential of these reefs. |
| **West coast of Grande Terre** | Reefs on the west coast of Grande Terre generally displayed higher levels of ICI and $PA_{HEAT}$, compared with the rest of the Archipelago. Under an adaptive potential perspective, reefs in the northern part are of paramount importance as they receive the propagules from all the south-western reefs that experienced frequent heat stress. No MPA is established in this area. Another strategic region are the reefs in front of Noumea, in the southern part of the west coast, since they were predicted with high $PA_{HEAT}$ and OCI. Here, more than 200 km<sup>2</sup> of protected areas are already established. |
| **South Lagoon** | The South Lagoon displayed heterogenous patterns of $PA_{HEAT}$ and connectivity. The highest $PA_{HEAT}$ were observed in the south-western extremity, which in turn was a region predicted with low OCI. The eastern part might be more interesting under a conservation perspective, as it was predicted with moderate $PA_{HEAT}$ and high OCI. These reefs are located upstream of the trade winds, and can simultaneously send propagules to both coasts of Grande Terre. A large marine reserve (180 km<sup>2</sup>) is already established to protect these reefs. As for the southern part of the east coast, coral nurseries of heat stress adapted colonies might increase the adaptive potential of this region. |
| **Northern reefs and Entrecasteaux reefs** | Northern reefs and Entrecasteaux reefs were predicted with moderate to high levels of $PA_{HEAT}$, and low values of OCI and ICI, compared with the reefs around Grande Terre. The critical region under an adaptive potential perspective might be the eastern part of Northern reefs. This is because these reefs depend on the incoming propagules from the east coast of Grande Terre, which are predicted with low $PA_{HEAT}$. |
| **Loyalty Islands, Astrolabe and Petri atolls** | The main conservation issue for all the reefs in this region is the low ICI. It is likely that arrival of propagules substantially depends on the reefs from Vanuatu (Fig. 1), located ~200 km upstream on the South Equatorial Current. Reefs in Ouvéa and Astrolabe atolls (already protected) might be of strategic importance, as they were predicted with moderate to high values of $PA_{HEAT}$ and OCI. Since reefs in Maré and Lifou showed low $PA_{HEAT}$, establishment of nurseries with heat stress adapted coral might be useful under an adaptive potential perspective. |

### Predictions on adaptive potential match coral cover

Heat exposure is considered to be one of the main drivers of coral mortality worldwide^11,31,32^. Our results were consistent with this view, as we found a significant negative association of coral cover with BAF_previous year_ (Fig. 5a). Adaptation might contribute to increase thermal tolerance in corals, but its potential depends on two elements: the existence of adapted corals and the presence of reef connectivity patterns facilitating their dispersal. In this study, we found both of these elements (PA_HEAT_ and ICI) as associated with reduced loss of coral cover after thermal stress.

Previous studies have reported reefs that display increased thermal tolerance after recurrent exposure to heat stress^7–11^, and recent research suggested that the thermal contrasts of New Caledonia might have driven adaptive processes in corals of the region^23^. Our results supported this view: while recent thermal stress (BAF_previous year_) was associated with a reduction in coral cover, this reduction was mitigated at reefs that have experienced past thermal stress and were therefore predicted with high PA_HEAT_ (Fig. 5e). In addition, PA_HEAT_ alone did not result in a significant association with coral cover rates (Fig. 5b), and this might be due to the fact that thermal adaptation is advantageous only in response to heat stress. Indeed, previous research reported trade-offs in traits involved in local adaptation and acclimatization to heat stress in corals ^33^. These trade-offs might explain why the highest rates of coral cover (>0.4) in absence of heat stress (BAF_previous year_=0) were mainly observed at reefs with low PA_HEAT_ (Fig. 5e).

Outbound connectivity was not found to be associated with changes in coral cover (Fig. 5c,f). This is not surprising, because beneficial effects of dispersal are expected at reefs receiving incoming propagules, rather than the opposite ^16,34^. Indeed, inbound connectivity was found to mitigate the negative association between BAF_previous year_ and coral cover (Fig. 5g). Two non-mutually exclusive reasons might explain this observation. First, high levels of incoming propagules might facilitate the turnover of dead colonies caused by heat stress^35^, although it has to be noted that this kind of recovery usually requires several years ^36^. Second, incoming dispersal facilitates the arrival of adapted propagules, and therefore promotes an adaptive response even at reefs that did not experience thermal stress before^37^. Indeed, we observed that the frequency of adaptive genotypes in *A. millepora* and *P. acuta* was generally higher at reefs predicted with low PA_HEAT_ and high ICI, than in those predicted with both low PA_HEAT_ and low ICI (Fig. S2). This view on genetic rescue via incoming migration is supported by the fact that every reef depends, to some extent, on its neighbors for larval recruitment^38^.

### Limitations and future directions

The associations found between changes in coral cover and the descriptors of thermal stress, probability of heat stress adaptation and connectivity do not necessarily imply causative relationships. Despite evidence of effects of thermal patterns on coral cover reported by previous studies, there might be other environmental constraints that are asymmetrical between the two coasts of Grande Terre and modulate coral cover changes. Further validation remains necessary and could be achieved via experimental assays of heat stress resistance^8^ in colonies sampled at reefs with different PA_HEAT_. This approach would also enable disentangling of the possible confounding role of acclimatization in heat stress adaptive responses ^12,33^.

Another important aspect to consider in future studies is the resolution of remote sensing datasets used for predictions. Here, we worked at a resolution of ∼5 km for thermal variables and ∼8.5 km for sea current data. While the overall environmental patterns appeared consistent with those characterized in previous studies, it is likely that small scale phenomena were neglected. For instance, reef heat stress exposure can vary substantially under the fine-scale (<1km) of a seascape^13^. The same applies to connectivity, since the use of high resolution (≤1 km) hydrodynamic models could improve the characterization of coral larvae fine-scale dispersal^39,40^.

A third limitation of our approach concerns the generalization of the biological and ecological characteristics of a reef. Here we assumed that the reef system of New Caledonia was a single homogenous ecological niche, hosting an “average” species with an “average” heat stress adaptive response. This simplification is useful to portray an overall prediction, but might lead to local inaccuracies. This is because the reef types of New Caledonia are variegated and species distributions varies accordingly^41,42^. Furthermore, different species have different levels of bleaching sensitivity^43^ and reproduce under different strategies^44^. For instance, the propagules of a broadcast spawning coral as *A. millepora* travel over longer distances, compared with those of brooding species as *P. damicornis* and *P. acuta*^45^. Consequently, *A. millepora* showed a lower rate of decrease of genetic correlation between corals per unit of seascape cost distance, in comparison with the *Pocillopora* species (Fig. 3).

In future studies, PA_HEAT_ and connectivity predictions should be calibrated to match these biological differences. It is for this reason that seascape genomics studies will become of paramount importance into the future, as they provide species-specific indications on 1) how thermal stress might be translated in probability adaptation, and 2) the biological meaning (e.g. degree of genetic separation) of a cost distance by sea currents^17,18^.

### Conclusions

In this study, we combined remote sensing of environmental conditions with genomic data to predict spatial patterns of heat stress adaptation and connectivity for the coral reefs of New Caledonia. We then retrieved field survey data and showed that recent heat stress was associated with a decrease in living coral cover, but also that such association appeared to be mitigated at reefs predicted with 1) high probability of heat stress adaptation and 2) high levels of incoming dispersal. The metrics computed in this work resumes the adaptive potential of corals against heat stress, and therefore represents valuable indices to support spatial planning of reef conservation.

## Methods

### Remote sensing of sea surface temperature

Satellite data characterizing sea surface temperature (SST) were retrieved from a publicly available database (dataset: ESA SST CCI reprocessed sea surface temperature analyses)^46,47^. This dataset provides daily records of SST at a ∼5 km resolution from the years 1981 to 2017 across the whole study area (Fig. 1). The shapes of the reef of the region^48^ were transformed into a regular grid (1,284 cells with maximal size of 5×5 km), and for each reef cell we extracted the average temperature for every day of the observational period using QGIS software^49^. We performed calculations of heat stress patterns in the R environment using the *raster* package (v. 3.0)^50,51^. For each reef cell, patterns of heat stress were computed using the bleaching alert definition developed by the Coral Reef Watch briefly described hereafter^20^. For every day, we calculated the “hotspot value” as the difference between SST and the maximal monthly mean (MMM, usually the monthly average of February in New Caledonia). The hotspot value was retained only when SST exceeded the MMM by at least 1 °C. Next, for each day, we calculated the cumulated hotspot values over the previous 84 days (3 months), and if this sum is > 0, the day is flagged as being ‘under bleaching alert’. Finally, we computed the frequency of days under bleaching alert for every year (BAF_year_) from 1985 to 2017. For the preceding years (1981-1984), BAF_year_ was not calculated such to avoid bias caused by estimating MMM over a limited number of years. An overall measure of BAF (BAF_overall_) was calculated as the average of all the BAF_year_ from 1985 to 2017.

### Seascape connectivity graph

For the estimation of connectivity we applied a method based on spatial graphs previously employed to study coral reef connectivity^17^ and briefly outlined hereafter. We retrieved a publicly available dataset describing the eastward and northward surface water velocity (Global Ocean Physics Reanalysis)^46^. This dataset provided daily records at ∼8.5 km resolution from 1993 to 2017. Since this resolution can be inaccurate close to coastlines, we increased the resolution to 1 km using the “resample” function (“bilinear” method) of the *raster* R package, and used high resolution bathymetry data (100 m resolution^52^) to remove the sea velocity value from pixels located on land. We then used the R package *gdistance* (v. 1.2)^53^ to create a matrix describing the transition costs between each adjacent pixel in the study area. These costs were inversely proportional to the frequency of transition based on sea currents. This seascape connectivity graph was calculated as the shortest cost distances across this matrix between for each pair of the 1,284 reef cells. Of note, two least-cost-paths were calculated for each pair of reef cells, one for each direction of the transition.

### Connectivity indices

The seascape connectivity graph was used to compute two indices connectivity for every reef cell of the study area: inbound connectivity and outbound connectivity. These indices had been defined in previous work on corals^17^ and were calculated in the R environment.

*- Outbound connectivity index (OCI)*: represents the predisposition of a reef to send coral propagules to its neighbors. For a given reef cell, it is calculated by defining all the neighboring reef cells that can be reached under a determined cost distance threshold (CDt). OCI is the total area (in km^2^) of the destination reef cells.

*- Inbound connectivity index (ICI)*: represents the predisposition of a reef to receive coral recruits from its neighbors. For a given reef cell, it is calculated by defining all the neighboring reef cells that can reach this target reef cell under a determined CDt. ICI is the total area (in km^2^) of these departure reef cells.

We set the value of CDt to 800 units in order to maximize the neighborhood without causing border effects. This value was calculated based on the reef cells’ cost distance to and from the borders of the study area (located ∼250 km around the most peripheral reef cells), where the minimal cost distances to and from the border were 836 and 801 units, respectively.

### SNPs dataset

We retrieved genomic data employed in previous seascape genomics analyses on three coral species of New Caledonia: *Acropora millepora, Pocillopora damicornis* and *Pocillopora acuta*^23^. This dataset encompassed more than one hundred individuals per population (167 in *A. millepora*, 118 in *P. damicornis*, 110 in *P. acuta*), collected at multiple sampling sites around Grande Terre (20 sites for *A. millepora*, 17 for *P. damicornis*, 17 for *P. acuta*) and genotyped using a Genotype-By-Sequencing approach^54^ characterizing thousands of single-nucleotide-polymorphisms (SNPs; 11,935 in *A. millepora*, 7,895 in *P. damicornis* and 8,343 in *P. acuta*). Of note, SNPs in this dataset were already filtered for rare allelic variants (minor allele frequency<0.05%) and linkage disequilibrium (LD-pruning threshold=0.3^55^).

### Probability of heat stress adaptation

The previous seascape genomics study investigated the genotype-environment associations between SNPs and 47 environmental descriptors (among which is BAF_overall_) using LFMM software^23,56^. In each of the three species, the analysis reported significant associations (q<0.01) of BAF_overall_ with potentially adaptive SNPs (10 in *A. millepora*, 18 in *P. damicornis*, and 4 in *P. acuta*). We employed these genotype-environment associations to predict the probability of heat stress adaptation (PA_HEAT_) from BAF_overall_ values. We used a method based on logistic regressions^21,57^ that was previously applied to corals^17^, with some modifications outlined hereafter.

For each individual used in the analysis, we retrieved the BAF_overall_ value at the sampling location. Next, we encoded the presence/absence of the putatively adaptive genotype as a binary variable using a custom function in the R environment. We then employed a generalize linear mixed model (GLMM) to evaluate how the presence/absence of putative adaptive genotypes (response variable) responded to BAF_overall_ (explanatory variable) across all the selected SNPs, individuals and species combined. This was done through the R package glmmTMB (v 1.0)^58^, using a logistic regression model where SNP identifier, sample identifier and species were introduced as random factor. The resulting model then was used to transform BAF_overall_ values associated with each of the 2,284 reef cells of New Caledonia in PA_HEAT_. The model was plotted using the visreg R package (v. 2.6.1)^59^.

### Reef connectivity and genetic correlations between corals

The SNPs dataset was used to evaluate whether reef connectivity predictions were representative proxies of the structure of three coral populations. In the R environment, we applied the following framework to the genotype matrix of each of the three species. First, we evaluated the relatedness between each pair of individuals in the dataset (13,861 pairs in *A. millepora*, 6,903 pairs in *P. damicornis*, 5,995 pairs in *P. acuta*) by calculating the genetic correlation (Pearson) based on SNPs values^60,61^. We then computed the distribution of the genetic correlation values, and excluded pairs of individuals with anomalously high or low correlation (*i*.*e*. exceeding the boundaries defined by the median of the distribution ± three times the interquartile range).

Next, we investigated the drivers of genetic correlations by using GLMMs designed through the R package glmmTMB^58^. We set two fixed effects as possible drivers of genetic correlation between individuals: ancestral distance and reef connectivity. Accounting for ancestral genetic structure is particularly important as corals are prone to hybridization or cryptic speciation^62,63^. The computation of ancestral distance featured the R package ALStructure (v. 0.1)^64^. For a given SNP matrix, ALStructure predicts the number of ancestral populations, and then estimates, for every individual, the admixture proportions to the ancestral populations.

For every pair of individuals, we then calculated the ancestral distance as the Euclidean distance between the respective admixture proportions. For which concerns reef connectivity, the fixed effect corresponded to the least-cost-path (from the seascape graph) linking the sampling sites of every pair of individuals. Since genetic correlations could not exceed the 0-1 boundaries, GLMMs were built using a beta regression^65^. The random factors in the GLMM were the identifiers of the individuals in the pairs, as well as the identifier of the pair of sampling sites.

Finally, we evaluated the relationship between ancestral distance, reef connectivity and genetic correlations of corals by 1) reporting the estimate and its standard deviation, as well as the p-value associated with Wald statistic^58^; 2) plotting the association using the visreg R package^59^.

### Coral cover data

Living coral cover data was retrieved from the 2017-18 report of the New Caledonian observational network of coral reefs (‘Réseau d’observation des récifs coralliens de Nouvelle Calédonie’, RORC)^24^. Overall, we used data from 74 survey stations distributed across the Archipelago of New Caledonia (Fig. 1). At each station, yearly coral cover surveys were performed along the same 100m transect using the “point intercept” technique. Surveys covered the period from 2003 to 2017, where 18 sites have been visited for less than five years, 27 for five to ten years, and 29 for more than ten years. The exact coordinates of survey stations were retrieved from the geographic information web-portal of New Caledonia (https://georep.nc/).

### Environmental characterization of survey sites

The coordinates of survey stations were used to find the corresponding reef cells and the associated values of the connectivity indices (OCI and ICI). For each survey record (i.e. survey at a given station in a specific year) we also calculated BAF_overall_ as the average BAF since 1985 to the year proceeding the survey. Based on the values of BAF_overall_ we computed PA_HEAT_ for each survey record. In addition, we calculated BAF values on a rolling temporal window describing average BAF for the year (BAF_previous year_) that preceded the year of survey.

### Analysis of coral cover change

We investigated the association of BAF_previous year_, PA_HEAT_ and connectivity indices (ICI and OCI; in total 4 explanatory variables) with coral cover rates (response variable) using GLMMs. This analysis focused on the coral cover rates of every survey record (total of 574 records). The computation of GLMMs was performed using the R package glmmTMB (v 1.0)^58^, which allowed us to model coral cover rates via beta regression ^65^. We accounted for the non-independence of survey records originated at the same station but on different years by setting the station effect as random factor on the coral cover rate^66^. This approach is recommended for studies of longitudinal data with irregular time points^67^. To avoid bias due to scale differences between explanatory variables, each variable was standardized to mean 0 and standard deviation 1 using the R “scale” function.

We built two types of GLMMs: univariate and bivariate. In univariate GLMMs, BAF_previous year_, PA_HEAT_, ICI and OCI were employed each as unique fixed effect. The goal was to determine whether the explanatory variables showed a standalone association (*i*.*e*. independent from other variables) with coral cover change. In bivariate models, GLMMs were constructed each with two fixed effects: 1) BAF_previous year_ and 2) the interaction between BAF_previous year_ and each of the remaining explanatory variables: PA_HEAT_, ICI and OCI. The goal of bivariate models was to investigate whether the potential effect of recent thermal stress (BAF_previous year_) on coral cover might be modulated by PA_HEAT_, ICI or OCI.

For each GLMM, we reported the estimate and its standard deviation, as well as the p-value (deemed significant when <0.05) associated with Wald statistic^58^ of the fixed effects. In addition, we compared the quality-of-fit of models by calculating the Akaike Information Criterion (AIC)^68^.

## Acknowledgments

We thank the *Réseau d’observation des récifs coralliens* of New Caledonia for collecting and sharing the field survey data. We thank Annie Guillaume for the comments and suggestions provided during the redaction of this paper. This work was supported by the United Nations Environment Programme (UNEP) and International Coral Reef Initiative (ICRI) coral reefs small grants programme (grant number: SSFA/18/MCE/005). We also thank the Government of France and the Government of the Principality of Monaco who provided the funding for the small grants.

