## supplementary figures for "Coral cover surveys corroborate predictions on reef adaptive potential to thermal stress"

**Supplementary Figure 1. Ancestral distance and genetic correlation between corals.** The three plots display genetic correlations between pairs of corals sampled in New Caledonia as a function of their ancestral distance (blue line, with the grey band showing the 95% interval of confidence). Genetic correlations were computed as the correlation of single-nucleotide-polymorphisms, while ancestral distance is the difference in admixture from the ancestral populations. Each plot displays this association for a different species (a: *Acropora millepora*, b: *Pocillopora damicornis*, c: *Pocillopora acuta*).

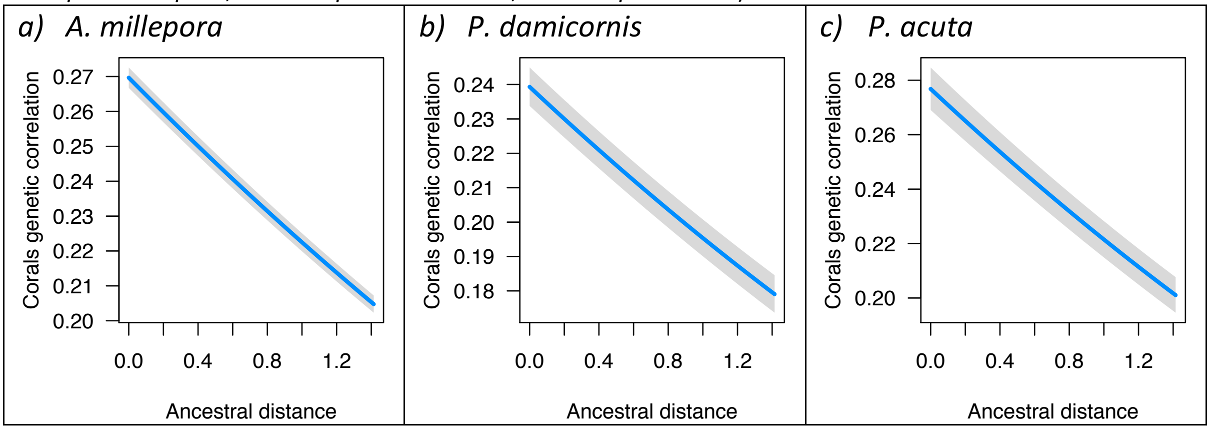


**Supplementary Figure 2. Observed frequency of genotypes adapted to heat.** The three boxplots display, for each of the three studied species (a: *Acropora millepora*, b: *Pocillopora damicornis*, c: *Pocillopora acuta*), the observed frequency of genotypes putatively adaptive to heat stress (y-axis) at reefs predicted with different values of probability of heat stress adaptation (low PA:PA<0.6, high PA: PA≥0.6) and inbound connectivity index (low ICI: ICI<1500 km$^2$, high ICI: ICI≥1500 km$^2$).

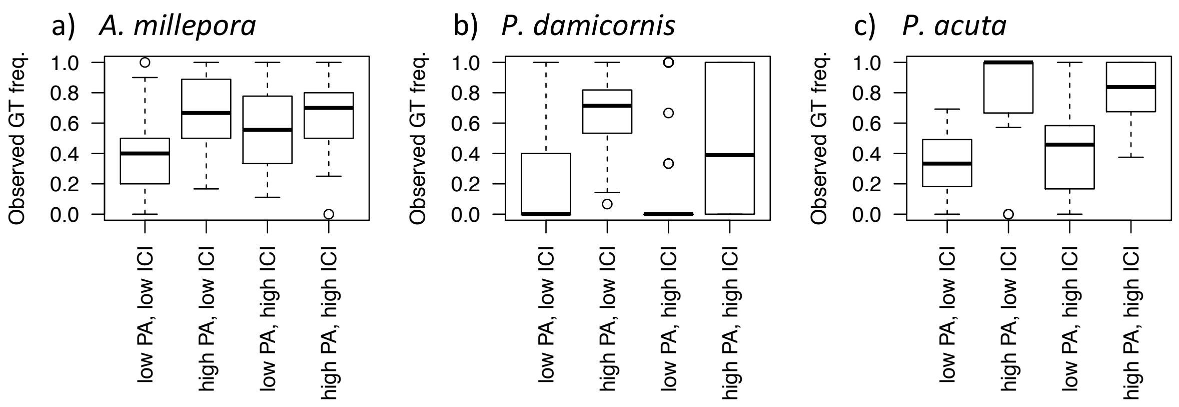
